# Suppression of PCBP1 Enhances PPARγ via TAK1 Modulation to Improve Glycemic and Lipid Metabolism Disorders in Gestational Diabetes Mellitus

**DOI:** 10.1101/2024.12.28.630638

**Authors:** Xuemei Xia, Yan Chen

**Affiliations:** Department of Obstetrical, Lishui People’s Hospital, Lishui 323000, China

**Keywords:** GDM, PCBP1, glucose and lipid metabolism, IR

## Abstract

Gestational diabetes mellitus (GDM) affects the health of pregnant women and their fetuses. Poly(C)-binding protein 1 (PCBP1), a multifunctional RNA-binding protein, is pivotal in maintaining cytosolic iron homeostasis. This study sheds light on the role and mechanism of PCBP1 in glucose and lipid metabolism dysregulation in GDM via a high-fat diet-induced GDM mouse model and a palmitic acid (PA)-triggered insulin resistance(IR) model in HepG2 cells. Glucose tolerance and insulin tolerance tests were performed in GDM mice, with lipid metabolism evaluated via hematoxylin-eosin (HE) staining, Oil Red O staining, as well as biochemical assay kits. Glucose content was quantified using the glucose oxidase method, while cell viability was evaluated trough the CCK-8 assay. Apoptotic activity was examined through TUNEL staining, and the expression levels of key proteins, including phosphorylated AKT (p-AKT), phosphorylated IRS1 (p-IRS1), PCBP1, TAK1, and PPARγ, were analyzed through Western blotting. The results demonstrated that GDM mice exhibited profound glucose and lipid metabolism disorders, characterized by significant lipid droplet accumulation in hepatic cells and disrupted insulin signaling pathways. Furthermore, hepatic expression of PCBP1 and TAK1 was notably upregulated, whereas PPARγ expression was significantly reduced. In vitro experiments revealed that silencing PCBP1 alleviated glucose and lipid metabolism abnormalities and improved insulin signaling in PA-induced insulin-resistant HepG2 (IR-HepG2) cells. This intervention also enhanced cell viability and suppressed apoptosis. Further mechanistic studies indicated that inhibition of TAK1 expression facilitated PPARγ upregulation, while TAK1 overexpression negated these effects. Additionally, silencing PPARγ in TAK1-silenced cells reversed the metabolic improvements in IR-HepG2 cells, whereas overexpression of PPARγ mitigated the adverse effects of PCBP1 overexpression. The foregoing findings demonstrate that PCBP1 exerts its effects on glucose and lipid metabolism in GDM via the TAK1/PPARγ signaling axis. Our study highlights the important function of the PCBP1/TAK1/PPARγ signaling pathway in mediating glucose and lipid metabolism in GDM, providing valuable insights into possible therapeutic targets for GDM treatment.

## Introduction

Gestational diabetes mellitus (GDM) is a disease featuring glucose metabolism abnormalities that manifest or are first identified during pregnancy [1]. It is linked to a risen risk of adverse pregnancy outcomes like embryonic loss, preeclampsia, dystocia, preterm birth, neonatal respiratory distress syndrome, and neonatal hypoglycemia. Furthermore, GDM increases the likelihood of developing diabetes, metabolic syndrome, and cardiovascular diseases in the postpartum period and later stages of life [2, 3]. At present, aside from lifestyle interventions and limited insulin therapy, no specific treatments for GDM have been established [4].

The pathogenesis of GDM is multifaceted, with an incidence rate exceeding 15% and continuing to rise. The onset of GDM is strongly associated with insulin resistance (IR) and pancreatic β-cell dysfunction, with dysregulated glucose and lipid metabolism serving as a key pathological hallmark [5, 6]. As pregnancy progresses, the placenta increasingly secretes insulin-resistant hormones, such as prolactin, progesterone, estradiol, and placental growth hormone, thereby escalating the demand for insulin production by pancreatic β-cells [7]. In normal pregnancies, β-cell proliferation and insulin secretion physiologically increase to counterbalance the heightened IR [8].

However, an abnormal elevation of insulin-resistant hormones and/or β-cell dysfunction in pregnant women can lead to a relative insulin deficiency, ultimately resulting in glucose and lipid metabolic dysregulation and the development of GDM [4]. Liver is pivotal in glucose and lipid metabolism by mediating gluconeogenesis, glycogenolysis, and lipid metabolic pathways to maintain systemic homeostasis. Hepatic dysfunction can disrupt glucose and lipid metabolism, thereby aggravating the progression of GDM [9]. Therefore, ameliorating IR and correcting glucose and lipid metabolic imbalances have become central objectives in the pursuit of effective GDM therapies.

Poly(C)-binding protein 1 (PCBP1) is a multifunctional RNA-binding protein involved in gene transcription, RNA regulation, and iron loading in ferritin [10]. Its expression is enriched in specific organ microenvironments and is dynamically modulated by the cellular environment, influencing cell growth, apoptosis, proliferation, differentiation and other processes[11, 12]. Skeletal muscle, a principal site of glucose uptake under insulin stimulation, relies on PCBP1 for the regulation of muscle differentiation via microRNA processing in myocytes [13, 14]. Studies have demonstrated that in rats with amyotrophic lateral sclerosis (ALS), impaired insulin signaling is accompanied by increased PCBP1 expression in muscle tissue [15]. Nevertheless, PCBP1’s involvement in the etiology of GDM is not elucidated.

The present study aims to unravel the effects of PCBP1 inhibition on glucose and lipid metabolic disorders associated with GDM, employing *in vivo* and *in vitro* models. Furthermore, our investigation uncovers the potential underlying mechanisms, thereby laying the theoretical groundwork for developing novel treatment strategies for GDM.

## 2. Materials and Methods

### 2.1. Cell culture and treatment

HepG2 cells (ATCC Cell Bank, China) were cultivated at 37°C in an atmosphere with 5% CO_2_ and saturated humidity in Dulbecco’s Modified Eagle Medium (DMEM) (11965092, Gibco, USA), supplemented with 10% fetal bovine serum (FBS) (22323002, Corning®, USA) and 1% penicillin-streptomycin (P7630, Solarbio, China).

The cells were treated to 0.25 mmol/L palmitic acid (PA) (Sigma, USA) for 24 hours in order to induce dysregulation of glucose and lipid metabolism.

### 2.2. Cell transfection

The logarithmic growth phase HepG2 and IR-HepG2 cells were planted in 12-well plates at a density of 1×10⁵ cells/mL. Following the manufacturer’s instructions, cell transfection was carried out using the Lipofectamine 2000 reagent (Thermo Fisher Scientific, USA). Plasmids such as si-NC, oe-vector, si-PCBP1, oe-PCBP1, si-TAK1, oe-TAK1, si-PPARγ, and oe-PPARγ (Shanghai GenePharma Co., Ltd., China) were present in the transfection, either separately or in combination. Cell samples and supernatants were gathered for further research after a 24-hour transfection period.

### 2.3. GDM model establishment

Eight-week-old C57BL/6N mice were purchased from the Zhejiang Center of Laboratory Animals (Production License: SCXK (Zhe) 2024-0002; Use License: SYXK (Zhe) 2024-0008) and acclimated for one week under standardized conditions of 20–26°C temperature and 50–56% humidity in individually ventilated cages at the Hangzhou Medical College Laboratory Animal Center. Male and female mice were housed separately. Female mice were randomized to a control group, which received a standard CHOW diet (11.8% calories from fat), or a high-fat diet (HFD) group, receiving a diet with 45% calories derived from fat (Beijing Botai Hongda Biotechnology Co., Ltd., China). Weekly body weights were recorded.

The appearance of vaginal plugs the next morning, known as gestational day 0.5 (GD0.5), indicated that the mating was successful. Female mice were matched with male mice at a 2:1 ratio after a 6-week feeding program. Mice that were pregnant were kept separately. The Zhejiang Center of Laboratory Animals’ Institutional Animal Care and Use Committee gave its approval to the experimental protocol.

### 2.4. Blood glucose and insulin level assessment

Blood samples were gathered via tail vein puncture from pregnant mice in the fasting state at 08:00 on GD0.5, GD9.5, and GD18.5 for blood glucose and insulin level evaluation. A glucose meter and test strips were leveraged to measure blood glucose (Roche, Switzerland). Pregnant mice having fasting blood glucose (FBG) levels ≥ 11.1 mmol/L were diagnosed with GDM as per established criteria [16]. Serum insulin concentrations were determined through a mouse insulin detection kit (Shanghai Enzyme Linked Biotechnology Co., Ltd., China).

### 2.5. Glucose tolerance test (GTT) and insulin tolerance test (ITT)

In GTT, mice were fasted for 6 hours (08:00-14:00 h) before FBG were obtained with a glucose meter and test strips, and the baseline glucose value was noted at 0 min. An intraperitoneal injection of a 20% physiological glucose solution at a dose of 2 g/kg body weight was then given. After the injection, blood glucose levels were measured 15, 30, 60, 90, and 120 min later to construct a glucose curve and calculate the area under the curve (AUC). ITT was conducted 72 hours after the GTT, with mice again fasting for 6 hours (08:00-14:00 h). The baseline blood glucose was recorded at 0 min, followed by an intraperitoneal injection of insulin at a dose of 0.5 IU/kg body weight (Sigma, USA). After the injection, blood glucose levels were checked 15, 30, 60, 90, and 120 min later. Mice were put to sleep with 2% sodium pentobarbital (50 mg/kg) and killed by cervical dislocation following the experiment. Serum and liver tissue samples were harvested and preserved for further analyses.

### 2.6. Serum biochemical index detection

As per the manufacturer’s instructions for every reagent kit, serum biochemical indices were evaluated in pregnant mice. Triglycerides (TG; A110-1-1, Nanjing Jiancheng Bioengineering Institute, China), total cholesterol (TC; A111-1-1, Nanjing Jiancheng Bioengineering Institute, China), low-density lipoprotein cholesterol (LDL-C; A113-1-1, Nanjing Jiancheng Bioengineering Institute, China), and high-density lipoprotein cholesterol (HDL-C; A112-1-1, Nanjing Jiancheng Bioengineering Institute, China) were quantified through measurement of absorbance at a wavelength of 500 nm.

### 2.7. Hematoxylin-Eosin (HE) staining

The liver tissues were preserved in 10% formalin, embedded in paraffin, cut with a microtome to an 8 μm thickness, and then placed on microscope slides. After that, the sections were deparaffinized, rehydrated, and stained for 2 min with hematoxylin solution and 30 s with eosin. Representative stained sections from each group were examined and photographed using a light microscope.

### 2.8. Western blot

Radio-immunoprecipitation assay (RIPA) lysis buffer (Shanghai Beyotime Biotechnology Co., Ltd., China) was employed for total cellular protein extraction. The bicinchoninic acid (BCA) method (Shanghai Beyotime Biotechnology Co., Ltd., China) was utilized to measure the concentrations of proteins. SDS-PAGE helped to separate the proteins (40 μg each sample), and a wet transfer method was then adopted to transfer the proteins onto PVDF membranes (Merck, Germany). For one hour, the membranes were blocked with 5% nonfat milk at room temperature. They were then incubated with primary antibodies, such as PCBP1 (ab168377, Abcam, China), TAK1 (ab109526, Abcam, China), PPARγ (ab272718, Abcam, China), p-IRS1 (ab4776, Abcam, China), IRS1 (ab245314, Abcam, China), p-AKT (#9271, CST, Germany), AKT (#9272, CST, Germany), GAPDH (ab9484, Abcam, China), and β-actin (#4967, CST, Germany) at 4°C for the entire night.

### 2.9. Quantitative real-time polymerase chain reaction (qRT-PCR)

The TRIzol reagent was utilized for extracting total RNA, which was subsequently reverse-transcribed into complementary DNA (cDNA). A qRT-PCR kit (TaKaRa, Japan) was used to measure the expression levels of PCBP1, TAK1, and PPARγ, with GAPDH acting as the internal reference. The 2 ^− Δ Δ Ct^ technique was applied for calculating relative gene expression. Pre-denaturation at 95°C for 5 min, 35 cycles of denaturation at 96°C for 30 s, annealing at 55°C for 30 s, and extension at 72°C for 20 s comprised the cycling settings. Supplementary Table S1 presents specific primer sequence information.

### 2.10. Cell counting kit-8 (CCK-8) assay

The CCK-8 assay (Shanghai Beyotime Biotechnology Co., Ltd., China) was helped to evaluate cell viability. In triplicate, cells were planted in 96-well plates. After 10 μL of CCK-8 reagent was added to each well, the plates were incubated for 3 h at 37°C in the dark. A microplate reader enabled the recording of the absorbance at 450 nm.

### 2.11. Terminal deoxynucleotidyl transferase dUTP nick end labeling (TUNEL) assay

An in situ cell death detection kit (Roche, Switzerland) assisted in assessing apoptosis. Cells were permeabilized with 0.2% Triton X-100 after being fixed in 4% paraformaldehyde. TUNEL was leveraged to mark DNA fragmentation in apoptotic cells., resulting in brown staining. Apoptotic cells were quantified by randomly selecting five fields per slide under a microscope.

### 2.12. Glucose oxidase method

Glucose concentrations in cell culture supernatants were obtained via the glucose oxidase detection kit (ml024803, Shanghai Enzyme Linked Biotechnology Co., Ltd., China).100 μL of HRP-conjugated detection antibody was added to 50 microliters of each sample or standard, and the mixture was incubated for one hour at 37°C. After a wash, 50 μL of glucose oxidase working solutions A and B were added to each well. Fifty microliters of stop solution were added after 15 min of dark incubation. Within 15 min, absorbance was measured at 450 nm, and a standard curve was employed to determine the glucose concentrations.

### 2.13. Oil red O staining

After being fixed for 30 min in 10% neutral-buffered formalin, the cells were equilibrated for 5 min in 60% isopropanol. The cells were then stained for ten min using a 0.5% Oil Red O solution. After staining, the excess stain was removed using 60% isopropanol until the background appeared clear. The cells were then washed three times with distilled water. To enhance visualization, we counterstained the cells with hematoxylin for 1 min, followed by an additional wash with distilled water. Finally, glycerol gelatin was applied for mounting, and lipid accumulation was observed microscopically.

### 2.14. Statistical analysis

Data were processed with the help of GraphPad Prism 9.0. Measurement data are displayed as *x*±s. Comparisons across groups were performed through one-way analysis of variance (ANOVA), and pairwise comparisons were completed through the t-test. Statistical significance was signified as \**p*<0.05; \*\**p*<0.01; \*\*\**p*<0.001; \*\*\*\**p*<0.0001.

## 3. Results

### 3.1 Metabolic dysregulation of glucose and lipids was observed in GDM mice

A GDM mouse model was constructed based on HFD. Blood glucose and insulin were examined at gestational days (GD) 0.5, 9.5, and 18.5. Blood glucose levels in the HFD group exceeded 11.1 mmol/L at GD9.5 and GD18.5 (p < 0.001) along with a significant decrease in insulin levels (p < 0.001), confirming the successful establishment of the GDM model (Figures 1A, B). GTTs showed that after glucose was administered, blood glucose levels rose in all groups, but the HFD group’s increase was more noticeable than the control group’s, peaking at 30 min. (Figure 1C). ITTs demonstrated impaired glucose recovery in the HFD cohort following insulin injection, compared to the control group (Figure 1D). Lipid metabolism analysis in pregnant mice revealed significantly elevated levels of TG, TC and LDL in the HFD cohort in comparison to controls, whereas HDL levels were markedly reduced (p < 0.01, Figure 1E). These findings indicate dysregulated glucose and lipid metabolism in the GDM model. Histological analysis of liver tissue via HE staining revealed lipid droplets of varying sizes within hepatocytes in the HFD group (Figure 1F). Previous studies have reported that PPARγ significantly impacts lipid transport in GDM, with TAK1 acting as a core regulator of PPARγ, which in turn is regulated by PCBP [17, 18]. Therefore, the protein expression levels of PCBP1, TAK1, and PPARγ were further investigated. In contrast to the control, the HFD cohort had notably elevated PCBP1 levels(*p* < 0.001) and TAK1 (*p* < 0.0001), while PPARγ and IR-linked proteins (p-IRS1 and p-AKT) were markedly suppressed (*p* < 0.001) (Figure 1G).

**Figure 1.**
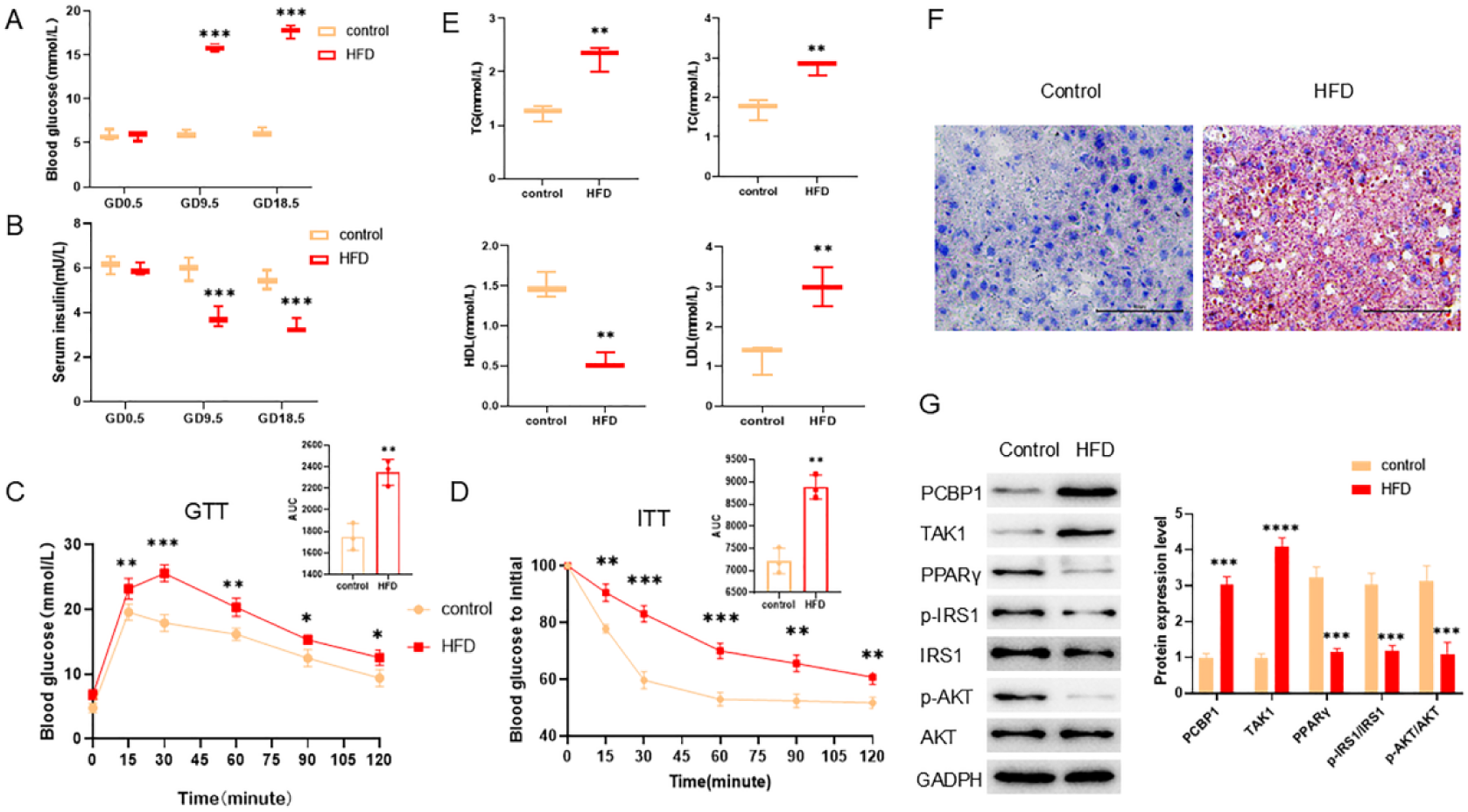
Metabolic dysregulation of glucose and lipids was observed in GDM mice. (A) Serum glucose levels in mice at GD 0.5, 9.5, and 18.5. (B) Serum insulin levels in mice at GD 0.5, 9.5, and 18.5. (C) Glucose tolerance of mice within 120 min. (D) Insulin tolerance of mice within 120 min. (E) Serum levels of TG, TC, HDL, and LDL. (F) HE staining of liver tissue. (G) The expression levels of PCBP1, TAK1, PPARγ, p-IRS1 (IRS1), and p-AKT (AKT) protein. \*\**p*<0.01; \*\*\**p*<0.001; \*\*\*\**p*<0.0001

### 3.2 Inhibition of PCBP1 improves glucose and lipid metabolism dysregulation and cell apoptosis in hepatocytes

The dysregulation of hepatic glucose and lipid metabolism constitutes a pivotal factor in the pathogenesis of GDM [19]. To elucidate the underlying mechanisms governing glucose and lipid metabolism, this study employed PA to treat HepG2 cells, thereby simulating the hepatic metabolic environment characteristic of women with GDM. Previous investigations have demonstrated that the inhibition of PCBP1 modulates the transcription of genes implicated in TG metabolism, transmembrane transport, and glucose metabolism in hepatocellular carcinoma cells [20]. Accordingly, a si-PCBP1 plasmid was designed to unveil the function of PCBP1 in glucose and lipid metabolic imbalance in HepG2 cells. In comparison to the si-NC group, the si-PCBP1 cohort exhibited marked silencing efficiency (*p* < 0.001) (Figure 1A). HepG2 cells treated with PA demonstrated reduced cell viability (p < 0.01), elevated glucose content (p < 0.0001), increased apoptosis, and augmented intracellular lipid droplet accumulation, confirming the successful establishment of the GDM-related cellular model. Following transfection with si-NC and si-PCBP1 plasmids, PA-treated HepG2 cells in the PA+si-PCBP1 group exhibited improved cell viability (*p < 0.05*), decreased glucose levels (*p < 0.0001*), reduced apoptosis, and a pronounced decline in intracellular lipid droplets relative to the PA+si-NC group (Figures 1B-E). These findings indicate that suppression of PCBP1 ameliorates glucose and lipid metabolic dysfunction, enhances cell viability, and mitigates apoptosis in HepG2 cells. Western blot analysis further showed significant changes of intracellular protein expression. Compared with the PA+si-NC group, the PA+si-PCBP1 cohort had less TAK1 expression (*p < 0.001*) and more PPARγ expression (*p < 0.05*). Additionally, the expression of IR-associated proteins p-IRS1 (*p < 0.001*) and p-AKT (*p < 0.01*) was markedly enhanced, indicating improved activation of insulin signaling pathways (Figure 1F).

### 3.3 Suppression of PCBP1 regulates glucose and lipid metabolism in hepatocytes via TAK1

Studies have reported that PCBP1 can bind to TAK1 precursor mRNA and exert its effects [21]. To investigate whether PCBP1 modulates PA-induced glucose and lipid metabolism in HepG2 cells via TAK1, our research first confirmed that the designed oe-TAK1 construct exhibited robust overexpression efficacy (*p < 0.0001*) (Figure 3A). HepG2 cells exposed to PA were co-transfected with either oe-vector or oe-TAK1 in combination with si-NC or si-PCBP1, or transfected individually. Notably, TAK1 overexpression mitigated the enhancements in cell viability (Figure 3B) (*p < 0.001*) and glucose uptake (Figure 3C) *(p < 0.001*) observed upon PCBP1 inhibition. Moreover, TAK1 overexpression exacerbated PA-induced cell apoptosis (Figure 3D) and intracellular lipid accumulation (Figure 3E). Western blot analyses revealed that PCBP1 inhibition significantly decreased endogenous TAK1 expression (p < 0.0001). Furthermore, the activation of IRS1 and AKT phosphorylation mediated by PCBP1 inhibition was reversed by TAK1 overexpression (*p < 0.05, p < 0.01*) (Figure 3F). Therefore, PCBP1 inhibition enhances insulin signaling by downregulating TAK1 expression, thereby ameliorating glucose and lipid metabolic dysfunction in HepG2 cells.

**Figure 2.**
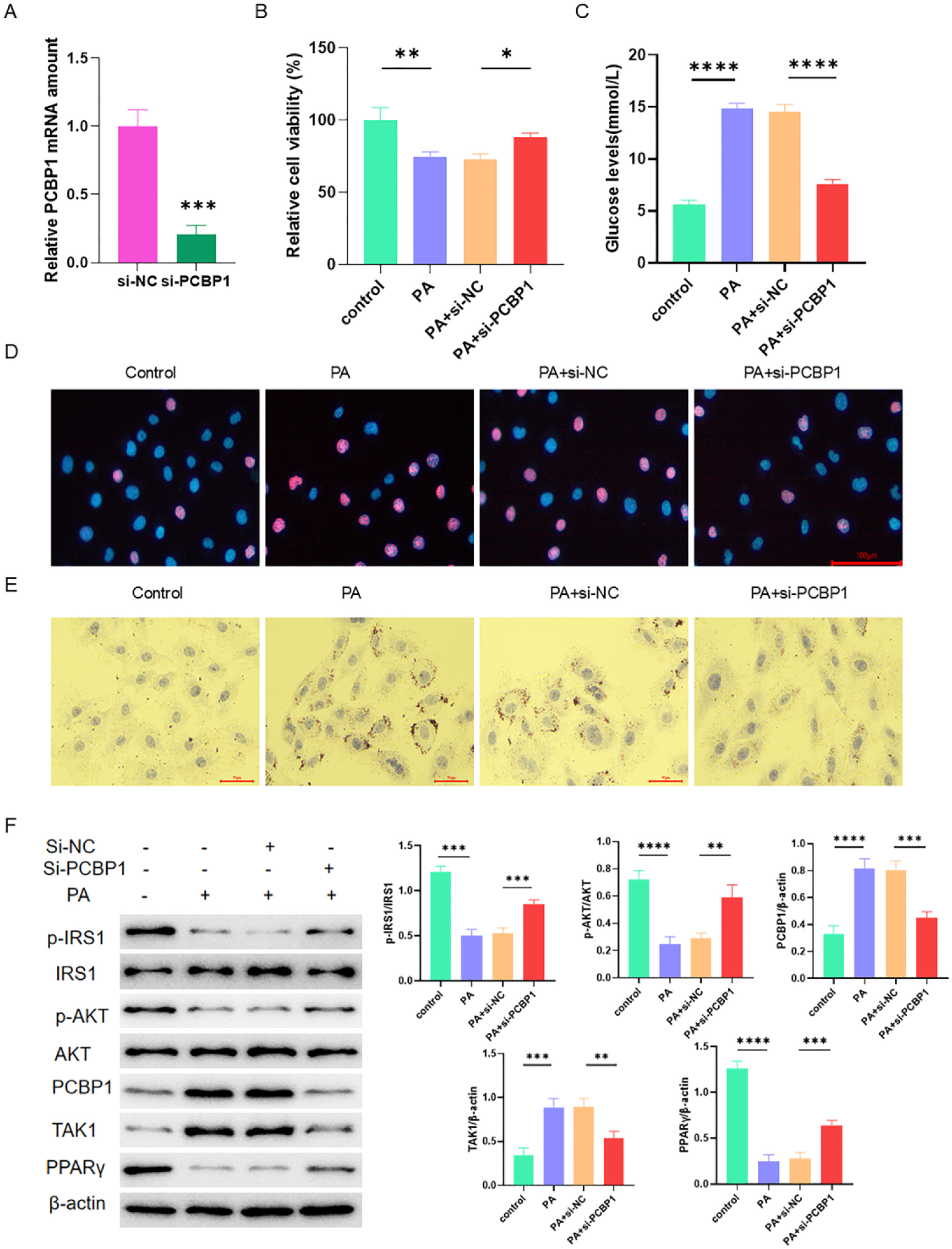
Inhibition of PCBP1 improves glucose and lipid metabolism dysregulation and cell apoptosis in hepatocytes. (A) Relative expression levels of PCBP1 mRNA (B) Cell viability were evaluated via the CCK-8 assay. (C) Glucose content in cells. (D) Cell apoptosis was measured by the TUNEL assay. Blue represents DAPI, and red refers to TUNEL. (E) Oil Red O staining showed cell morphology. (F) The expression levels of PCBP1, TAK1, PPARγ, p-IRS1 (IRS1), and p-AKT (AKT) protein. \**p*<0.05; \*\**p*<0.01; \*\*\**p*<0.001; \*\*\*\**p*<0.0001.

**Figure 3.**
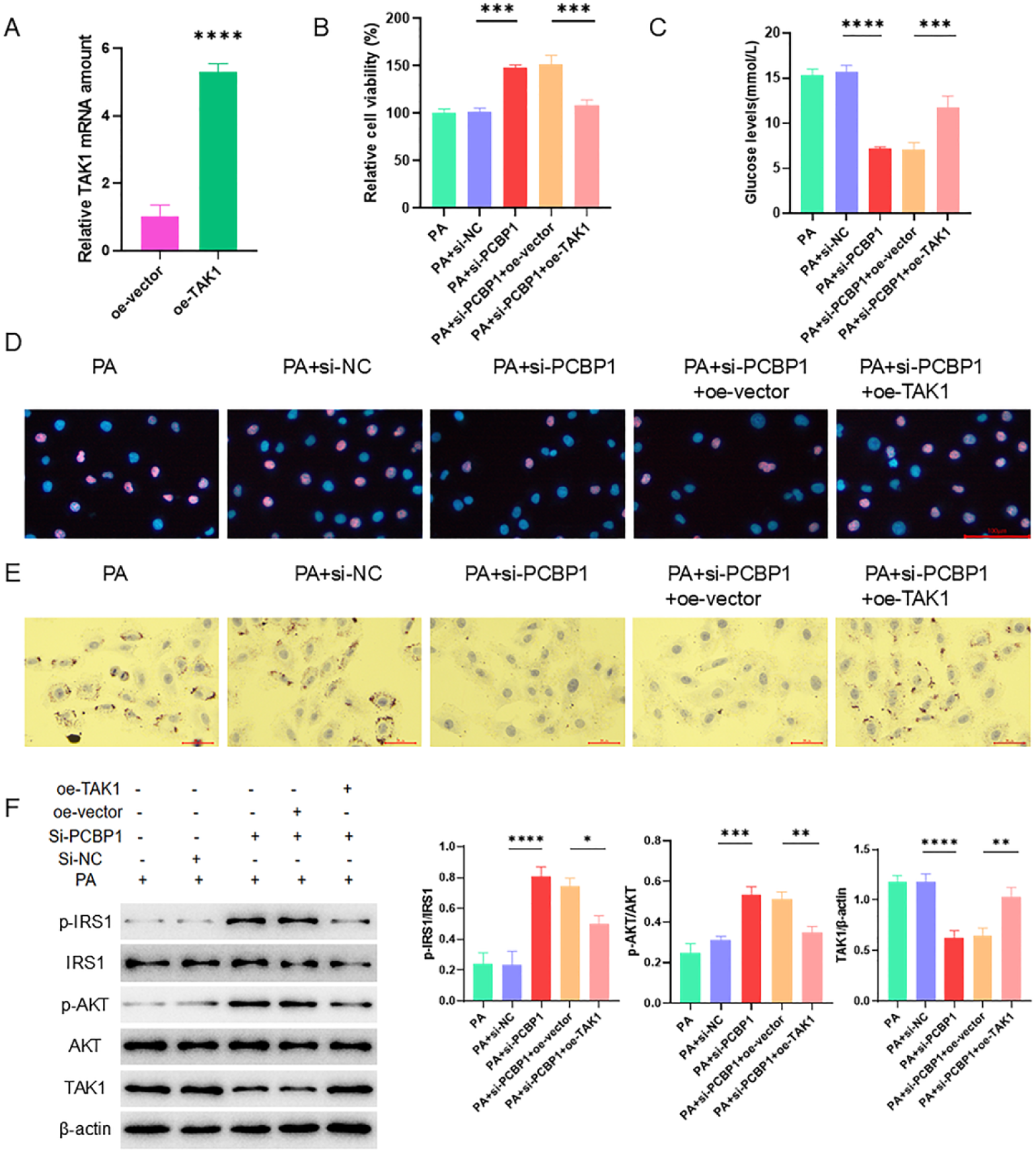
Inhibition of PCBP1 regulates glucose and lipid metabolism in hepatocytes via TAK1. (A) Relative expression levels of TAK1 mRNA. (B) Cell viability was assessed through the CCK-8 assay. (C) Glucose content in cells. (D) Cell apoptosis was measured via the TUNEL assay. Blue represents DAPI, and red refers to TUNEL. (E) Oil Red O staining was used to examine cell morphology. (F) The expression levels of PCBP1, TAK1, PPARγ, p-IRS1 (IRS1), and p-AKT (AKT) protein. \**p*<0.05; \*\**p*<0.01; \*\*\**p*<0.001; \*\*\*\**p*<0.0001.

### 3.4 TAK1 regulates glucose and lipid metabolism in hepatocytes via PPAR**γ**

TAK1 is a critical upstream molecule regulating the transcriptional activity of PPARγ [18]. This study further investigated whether TAK1 regulates glucose and lipid metabolism in PA-triggered HepG2 cells through PPARγ. First, si-PPARγ and si-TAK1 plasmids were designed, and significant silencing effects were confirmed (*p* < 0.001) (Figure 4A). Co-transfection or individual transfection of si-NC, si-PPARγ, and si-TAK1 into PA-induced HepG2 cells revealed that the increase in cell viability caused by silencing endogenous TAK1 expression was reversed by simultaneous PPARγ silencing (Figure 4B) (*p* < 0.001). Furthermore, silencing PPARγ expression exacerbated intracellular glucose production (Figure 4C) (*p* < 0.001), increased apoptosis (Figure 4D), and led to a further accumulation of intracellular lipid droplets (Figure 4E). To further verify the link of TAK1 to PPARγ, relevant protein expression was examined via Western blot analysis, which revealed that TAK1 silencing upregulated PPARγ expression and boosted the transcription of insulin-related proteins. However, when both TAK1 and PPARγ were silenced, transcription of p-IRS1 and p-AKT was inhibited. These findings suggest that PCB1’s downstream TAK1 may influence glucose and lipid metabolism in HepG2 cells by regulating PPARγ (Figure 4F).

**Figure 4.**
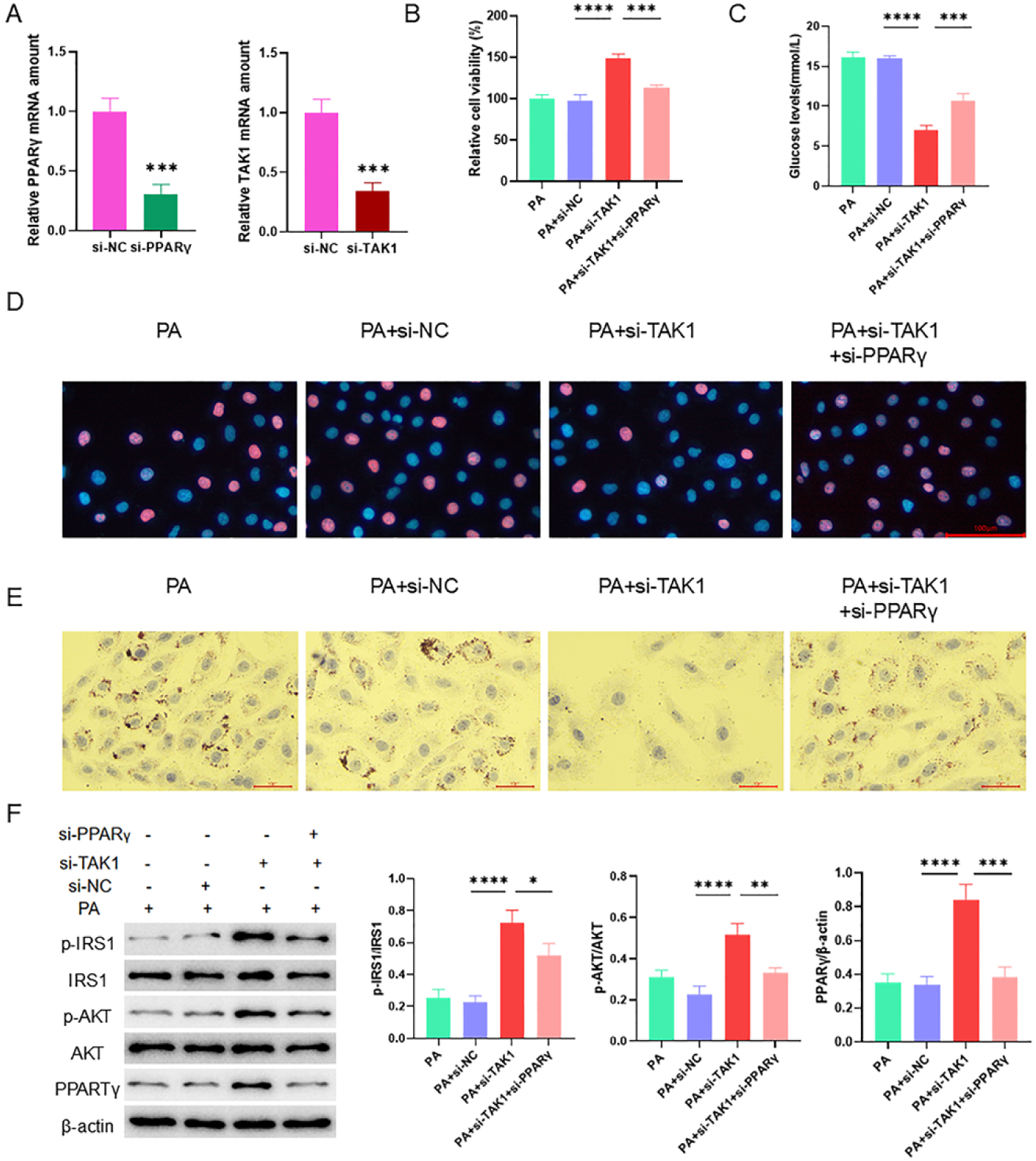
TAK1 regulates glucose and lipid metabolism in hepatocytes via PPARγ. (A) Relative expression levels of PPARγ and TAK1 mRNA. (B) Cell viability was evaluated using the CCK-8 assay. (C) Glucose content in cells. (D) Cell apoptosis was assessed by the TUNEL assay. Blue represents DAPI, and red refers to TUNEL. (E) Oil Red O staining showed cell morphology. (F) The expression levels of PPARγ, p-IRS1 (IRS1), and p-AKT (AKT) protein. \**p*<0.05; \*\**p*<0.01; \*\*\**p*<0.001; \*\*\*\**p*<0.0001.

### 3.5 PCBP1 regulates PPARγ expression via TAK1 to affect glucose and lipid metabolism in hepatocytes

To unveil the molecular mechanisms by which PCBP1 regulates glucose and lipid metabolism in hepatocytes, overexpression plasmids for PCBP1 (oe-PCBP1) and PPARγ (oe-PPARγ) were constructed, and their significant overexpression was validated (p < 0.001) (Figure 5A). Transfection of these plasmids, either individually or in combination, into PA-induced HepG2 cells demonstrated that PPARγ overexpression enhanced cell viability and glucose uptake, whereas simultaneous overexpression of PCBP1 significantly inhibited cell growth (p < 0.0001) and further promoted glucose accumulation in hepatocytes (p < 0.05) (Figures 5B and 5C). Furthermore, co-overexpression of PCBP1 increased cell apoptosis and intracellular lipid secretion (Figures 5D and 5E). At the protein expression level, PPARγ overexpression augmented the phosphorylation of IRS1 (p-IRS1) and AKT (p-AKT) without affecting the levels of PCBP1 and TAK1. However, concurrent overexpression of PCBP1 suppressed the expression of p-IRS1, p-AKT, and PPARγ while upregulating TAK1 expression (Figure 5F). Collectively, these findings, in conjunction with previous studies, suggest that inhibiting PCBP1 may enhance PPARγ expression by suppressing TAK1, thereby ameliorating glucose and lipid metabolic disorders in hepatocytes.

**Figure 5.**
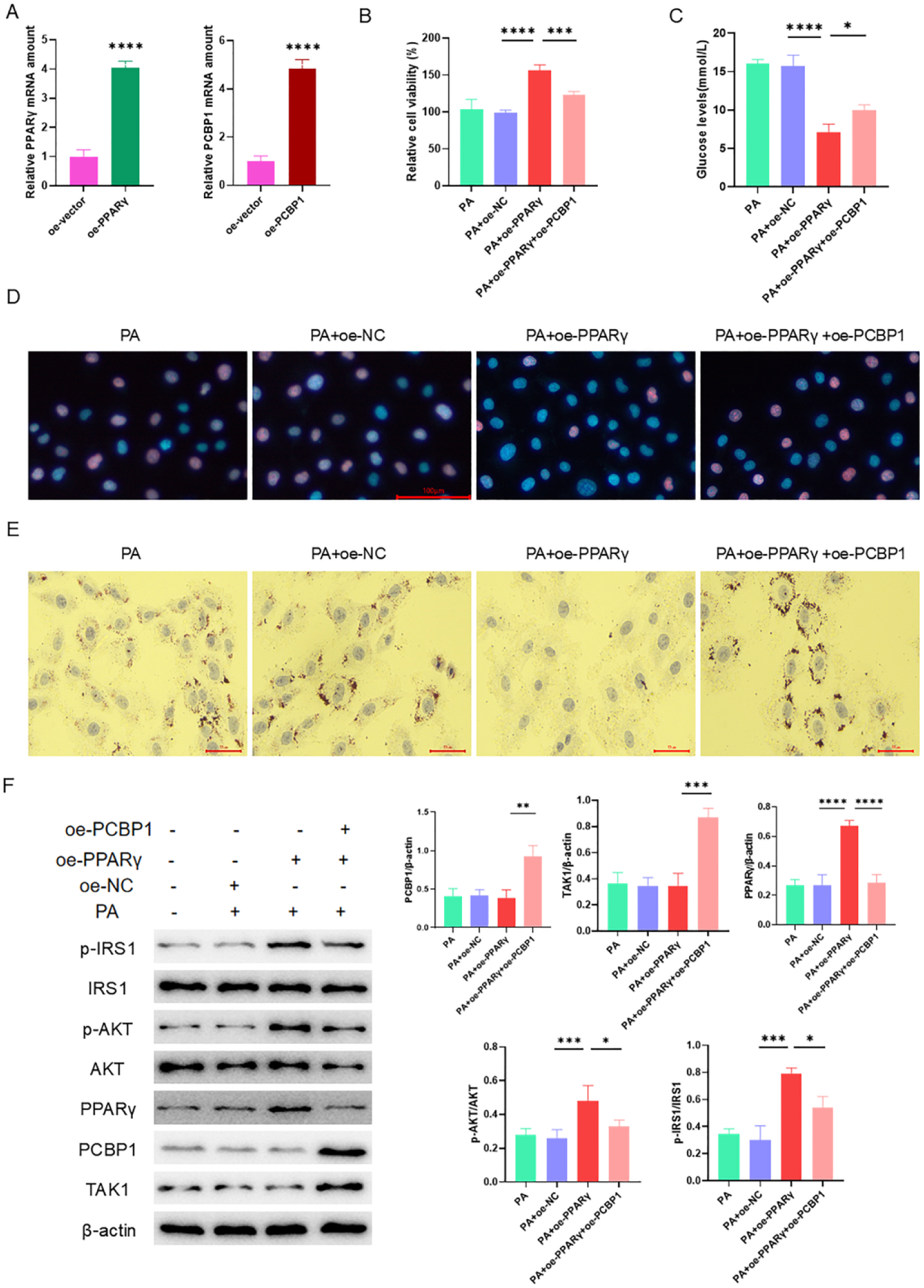
PCBP1 regulates PPARγ expression via TAK1 to affect glucose and lipid metabolism in hepatocytes. (A) Relative PPARγ and PCBP1 mRNA expression levels. (B) Cell viability was gauged using the CCK-8 assay. (C) Glucose content in cells. (D) Cell apoptosis was evaluated via the TUNEL assay. Blue represents DAPI, and red refers to TUNEL. (E) Oil Red O staining was used to examine cell morphology. (F) The expression levels of PCBP1, TAK1, PPARγ, p-IRS1 (IRS1), and p-AKT (AKT) protein. \**p*<0.05; \*\**p*<0.01; \*\*\**p*<0.001; \*\*\*\**p*<0.0001.

## 4 Discussion

This study aimed to unravel the impact of PCBP1 on glucose and lipid metabolism dysregulation in GDM and to explore its potential molecular mechanisms. The findings demonstrated that inhibition of PCBP1 expression significantly ameliorated glucose and lipid metabolism dysregulation by suppressing TAK1 expression and upregulating PPARγ. These results underscore the pivotal role of PCBP1 in maintaining glucose and lipid homeostasis and indicate that the PCBP1/TAK1/PPARγ signaling pathway is a promising therapeutic target for the treatment of GDM-associated metabolic abnormalities.

In recent decades, lifestyle and dietary shifts have contributed to a rise in obesity and related risk factors, resulting in an accelerated prevalence of GDM. Therefore, developing effective therapeutic strategies for GDM is pressing [22]. Previous studies have reported a strong relation of maternal glucose and lipid metabolism dysregulation tp the incidence of GDM [23]. In GDM, hepatic metabolic function is often profoundly altered [24, 25]. Consistent with these findings, our GDM mouse model exhibited persistent abnormalities in glucose uptake and insulin secretion. Furthermore, histological examination of liver tissues revealed pronounced steatosis and lipid droplet infiltration in the livers of GDM-affected pregnant mice, accompanied by elevated PCBP1 and TAK1 expression and reduced PPARγ expression.

PCBP1 belongs to a family of adaptor proteins interacting with poly(C)-rich RNA, DNA, and iron complexes and regulating cytosolic iron homeostasis in yeast and mammalian cells [26]. Previous studies have shown that PCBP1 deficiency can induce weight loss in mice and impair intestinal iron absorption [27]. Moreover, PCBP1 has been demonstrated to bind to the precursor mRNA of TAK1, a process enhanced by TGF-β signaling [21]. To further elucidate the effects of PCBP1 on GDM, our in vitro experiments revealed that inhibition of PCBP1 expression significantly improved glucose uptake, reduced lipid accumulation, and enhanced insulin signaling in IR-HepG2 cells. PCBP1 inhibition also decreased cell apoptosis and promoted cell viability, indicating its influence on the pathophysiology of GDM. We investigated the function of TAK1, a serine/threonine kinase that is essential for the pathways involved in inflammation and stress response, in order to clarify the molecular mechanisms by which PCBP1 modulates glucose and lipid metabolism [28]. Previous research has indicated that TAK1 degradation mediated by E3 ligase TRIM16 alleviates lipid accumulation and inflammation in a mouse model of non-alcoholic steatohepatitis (NASH) [29]. Additionally, suppression of TAK1 signaling can mitigate NASH progression induced by metabolic stress [30]. In the present study, overexpression of TAK1 abrogated the metabolic improvements resulting from PCBP1 downregulation, suggesting that PCBP1 exacerbates IR by activating the TAK1 pathway. These findings align with the low-grade chronic inflammation characteristic of GDM [31], further supporting the role of TAK1 as a mediator of PCBP1’s effects.

Our results further demonstrated that inhibition of PCBP1 upregulates PPARγ by suppressing TAK1 expression, thereby ameliorating glucose and lipid metabolism dysregulation in GDM. PPARγ, a ligand-dependent nuclear transcription factor, is a master mediator of adipocyte gene expression and insulin β-cell signaling. It is integral to glucose and lipid metabolism, immune response modulation, and cellular differentiation [32–34]. TAK1 has been identified as a critical upstream regulator of PPARγ transcriptional activity, interacting with TAK1-binding protein 1 (TAB1) to modulate PPARγ function and influence intracellular lipid synthesis [18, 35]. Dysfunction of PPARγ is related to many diseases like diabetes, atherosclerosis, and neurodegenerative disorders [36–38]. Furthermore, recent studies have reported that TAK1 inhibition activates PPARγ, potentially reducing cancer cell resistance and exerting antitumor effects [39]. Aligning with the foregoing findings, our study proved that PPARγ silencing reversed the metabolic benefits conferred by TAK1 inhibition, establishing PPARγ as a critical downstream effector in the TAK1-mediated regulation of glucose and lipid metabolism. Additionally, overexpression of PCBP1 exacerbated metabolic dysregulation, an effect reversed by increased PPARγ expression. The findings collectively indicate that the inhibition of PCBP1 mitigates glucose and lipid metabolism dysregulation via the downregulation of TAK1 and upregulation of PPARγ.

Thus, the PCBP1/TAK1/PPARγ signaling axis emerges as a potential therapeutic target for GDM. Although PPARγ agonists, such as thiazolidinediones, have been employed in type 2 diabetes treatment, their efficacy in GDM remains underexplored. This study highlights the potential of targeting the PCBP1/TAK1/PPARγ axis to address the therapeutic challenges associated with GDM [40].

In conclusion, our study elucidated the critical regulatory function of PCBP1 in the dysregulation of glucose and lipid metabolism in GDM, utilizing in vivo and in vitro experimental models. PCBP1 was shown to regulate insulin sensitivity and lipid metabolism through the TAK1/PPARγ signaling pathway. This regulatory axis provides a novel therapeutic target and theoretical framework for GDM treatment. Future investigations should further explore the clinical applicability of this pathway, offering new perspectives for individualized therapeutic strategies for GDM and advancing fundamental research into glucose and lipid metabolism disorders.

## Availability of data and material

The datasets used and/or analyzed during the current study are available from the corresponding author on reasonable request.

## Acknowledgment

We sincerely appreciate the investigators and authors who have contributed to this field. This research was supported by the Zhejiang Provincial Medical and Health Technology Project (Grant No. 2022498503).

## Author contributions

Xuemei Xia, analysis and collection of the data, drafting the manuscript; Yan Chen, conception and design, analysis and explanation of the data, revising the manuscript. All the authors have read and approved the final manuscript.

